# excluderanges: exclusion sets for T2T-CHM13, GRCm39, and other genome assemblies

**DOI:** 10.1101/2022.11.21.517407

**Authors:** Jonathan D. Ogata, Wancen Mu, Eric S. Davis, Bingjie Xue, J. Chuck Harrell, Nathan C. Sheffield, Douglas H. Phanstiel, Michael I. Love, Mikhail G. Dozmorov

## Abstract

**Summary:** Exclusion regions are sections of reference genomes with abnormal pileups of short sequencing reads. Removing reads overlapping them improves biological signal, and these benefits are most pronounced in differential analysis settings. Several labs created exclusion region sets, available primarily through ENCODE and Github. However, the variety of exclusion sets creates uncertainty which sets to use. Furthermore, gap regions (e.g., centromeres, telomeres, short arms) create additional considerations in generating exclusion sets. We generated exclusion sets for the latest human T2T-CHM13 and mouse GRCm39 genomes and systematically assembled and annotated these and other sets in the *excluderanges* R/Bioconductor data package, also accessible via the BEDbase.org API. The package provides unified access to 82 GenomicRanges objects covering six organisms, multiple genome assemblies and types of exclusion regions. For human hg38 genome assembly, we recommend *hg38.Kundaje.GRCh38_unified_blacklist* as the most well-curated and annotated, and sets generated by the Blacklist tool for other organisms.

**Availability and implementation:** https://bioconductor.org/packages/excluderanges/

**Contact:** Mikhail G. Dozmorov

**Supplementary information:** Package website: https://dozmorovlab.github.io/excluderanges/

## Introduction

Up to 87% of sequencing reads generated by chromatin targeting technologies (e.g., ChIP-seq) can map to a reference genome in distinct clusters (aka high-signal pileups)^1,2^ (*1*). These pileups frequently occur in regions near assembly gaps, copy number-high regions, and in low-complexity regions (*2*, *3*). Removing reads overlapping those regions, referred hereafter as exclusion sets, improves normalization of the signal between samples, correlation between replicates, and increases accuracy of both peak calling and differential ChIP-seq analysis (*4*–*6*). Therefore, standardized availability of those exclusion sets is critical for improving reproducibility and quality of bioinformatics analyses.

Finding and choosing an exclusion set can be a non-trivial task. The ENCODE project returns 94 hits using the “exclusion” search term (as of 11/08/2022)^3^, most of them having minimal annotation and unknown curation methods. These sets are available for human and mouse genome assemblies; however, the ENCODE project lacks exclusion sets for the latest Telomere-to-Telomere (T2T-CHM13) human and Genome Reference Consortium Mouse Build 39 (GRCm39/mm39) mouse assemblies. Converting exclusion set coordinates between genomic assemblies using liftOver is not advisable since new artifact-prone regions are added and others are lost due to closed gaps (*1*); therefore, exclusion sets should be generated and used for their respective genome assemblies. Furthermore, exclusion regions have been observed in genomes of other species and many exclusion sets for model organisms remain unpublished and scattered across GitHub repositories. We curated a collection of exclusion sets for six model organisms and 12 genome assemblies, including the newly generated T2T and mm39 exclusion sets. We included two other types of potentially problematic regions: University of California Santa Cruz (UCSC)-annotated gap sets, e.g., centromere, telomere, short arm, and Nuclear mitochondrial (NUMT) sets containing mitochondrial sequences present in the nuclear genome (*7*). We assemble a total of 82 uniformly processed and annotated exclusion sets in the *excluderanges* R/Bioconductor data package and provide API access via BEDbase.org.

## Implementation

An overview of the *excluderanges* data is shown in Figure 1. To create this resource, we performed a systematic internet and literature search. The ENCODE project was the largest source of exclusion sets for human (11 sets) and mouse (6 sets) organisms, covering hg19, hg38, mm9, and mm10 genome assemblies. We also obtained exclusion sets generated by the Blacklist (*1*) and PeakPass (*5*) software. Additionally, we obtained exclusion sets for *C. elegans* (ce10 and ce11 genome assemblies), *D. melanogaster* (dm3 and dm6), *D. rerio* (danRer10), and *A. thaliana* (TAIR10). Using the Blacklist software, we generated exclusion sets for the latest Telomere-to-Telomere (T2T-CHM13) human and Genome Reference Consortium Mouse Build 39 (GRCm39/mm39) mouse assemblies (Table 1, Supplementary Table S1).

**Figure 1.**
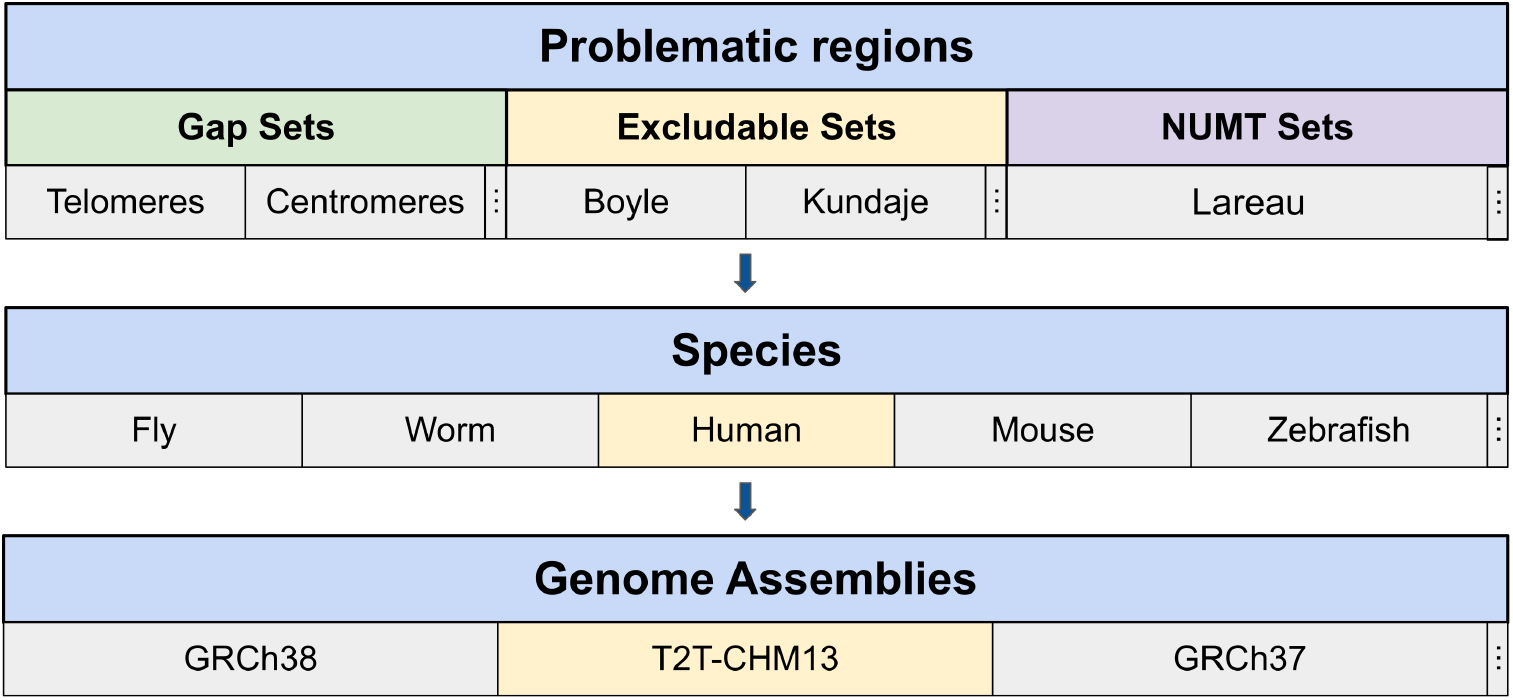
Schematic overview of the *excluderanges* package. Data for each type of problematic region (exclusion sets, gaps, Nuclear Mitochondrial (NUMT) sets) were obtained from public sources for each model organism and the corresponding genome assemblies. Exclusion sets for T2T-CHM13 and GRCm39 genome assemblies were *de novo* generated. Three vertical dots indicate more categories in the corresponding section.

**Table 1.** Characteristics of recommended exclusion sets for human and mouse genome assemblies. Unless specified otherwise, exclusion sets were defined by the Boyle-Lab/Blacklist software. The complete list is provided in Supplementary Table S1.

| Name | Assembly | Number of regions | Width, min/median/max, bp | Percent of the genome, % | Year last updated |
| --- | --- | --- | --- | --- | --- |
| *T2T.excluderanges* | T2T | 2066 | 1001/9701/25738901 | 8.358 | 2022 |
| *hg38.Kundaje.GRCh38_unified_Excludable*<sup>7</sup> | hg38 | 910 | 19/384/5407756 | 2.317 | 2020 |
| *hg38.Boyle.hg38-Excludable.v2* | hg38 | 636 | 1200/10150/30590100 | 7.355 | 2018 |
| *hg38.Wimberley.peakPass60Perc_sorted*<sup>8</sup> | hg38 | 5078 | 1000/2000/1852000 | 2.387 | 2021 |
| *hg19.Boyle.hg19-Excludable.v2* | hg19 | 834 | 1100/9350/30590100 | 8.882 | 2018 |
| *mm39.excluderanges* | mm39 | 3147 | 1100/12500/5487000 | 6.272 | 2022 |
| *mm10.Boyle.mm10-Excludable.v2* | mm10 | 3435 | 1000/8100/50585400 | 8.768 | 2018 |

Mitochondrial DNA sequences (mtDNA, 100-600K mitochondria per human cell) transferred to the nucleus give rise to the so-called mitochondrial DNA sequences in the nuclear genome (NUMTs). These sequences are found in genomes of various species (*7*), suggesting NUMTs may be a pervasive phenomenon. In the settings of DNA/chromatin sequencing (e.g., ATAC-seq), up to 80% of mitochondrial sequencing reads (*8*) may pile up in the NUMT sequences. Similar to exclusion sets, genomic regions highly homologous to mtDNA can be masked to improve biological signal. The reference human nuclear mitochondrial sequences have been available in the UCSC genome browser for hg18 (RHNumtS.2 database (*9*)) and lifted over to hg19 human genome assembly. Similarly, mouse NUMTs (RMNumtS database (*10*)) are available for the mm9 mouse genome assembly. However, recent human, mouse, and other organism genome assemblies lack NUMTs annotations in the UCSC database. We collected NUMT sets for more recent human and mouse genome assemblies, including hg38, T2T-CHM13, mm10, generated by Caleb Lareau in the mitoblacklist GitHub repository^4^.

Gaps in the genome represent another type of problematic regions. These include centromere and telomere sequences, short arms, gaps from large heterochromatin blocks, etc. While some are present in genome assemblies of most organisms (centromeres, telomeres, short arms, covering 2.47% ± 1.64, 0.01% ± 0.01, and 15.39%±3.66 of hg38 chromosomes, respectively), many are assembly-specific (e.g., gaps between clones, contigs, scaffolds in hg19 and hg38 assemblies). Gap data are available from the UCSC Genome Browser database or UCSC-hosted data hubs. The T2T-CHM13 assembly lacks assembly-specific gaps by the definition of telomere-to-telomere sequencing (*11*); however, coordinates of centromeres and telomeres are available from the CHM13 GitHub repository^5^. Additionally, we obtained T2T peri/centromeric satellite annotations, known to be associated with constitutive heterochromatin and span sites involved in kinetochore assembly or sequences epigenetically marked as centromeres (*12*). We also included the rDNA gap regions and regions unique to T2T-CHM13 v2.0 as compared with GRCh38/hg38 and GRCh37/hg19 assemblies under the rationale that alignments within these previously problematic regions might warrant extra attention. We characterized hg38 exclusion sets for overlap with gap regions and found that *hg38.Kundaje.GRCh38_unified_Excludable, hg38.Boyle.hg38-Excludable.v2*, and *hg28.Wimberley.peakPass60Perc_sorted* cover 99.40%, 99.08%, and 59.60% of centromeric regions, respectively. Notably, relatively few large regions were responsible for these overlaps (e.g., 27 out of 910 in *hg38.Kundaje.GRCh38_unified_Excludable*). In contrast, over 60% of the *hg38.Nordin.CandRblacklist_hg38* exclusion set for the CUT&RUN technology overlapped centromeres on chromosomes 1 and 13. Only sets generated by the Blacklist software overlapped centromeres, telomeres, and short arms, and there results were consistent across organisms and genome assemblies (Supplementary Table S2). Given the distinct properties of gap regions and inconsistency of their presence in exclusion sets, the aforementioned NUMTs and gap sets may be combined with other exclusion sets.

The large number of exclusion sets (e.g., nine for hg38 human genome assemblies) creates uncertainty in which set to use for a given genome assembly. We annotated exclusion sets by their creation methods, date of last update, width distribution, percent of the genome covered, and other properties (Supplementary Table S1, BEDbase.org^6^). Only sets generated by the Boyle’s lab Blacklist (*1*) or PeakPass by Eric Wimberley (*5*) software had published methods. While methods for some sets may be inferred (e.g., the hg38 *Yeo.eCLIP_Excludableregions.hg38liftover* set may have been lifted over from hg19), we advise against using poorly annotated sets. We also characterized hg38 exclusion sets and found they vary dramatically in terms of number (12,052 - 38) and width (median 10,151 - 30bp) (Supplementary Figure S1A, B). We calculated Jaccard overlap between each pair of hg38 exclusion sets, 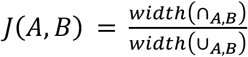. We found that *hg38.Kundaje.GRCh38_unified_Excludable* had the best Jaccard overlap with other sets, followed by_hg38.Wimberley.peakPass60Perc_sorted_ and *hg38.Boyle.hg38-Excludable.v2* sets (Supplementary Figure S1C). We additionally calculated overlap coefficient 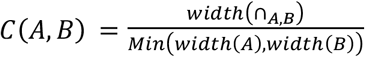 to minimize the effect of set size differences. We similarly found Kindaje-generated sets showing the best overlap with other sets, followed by *hg38.Boyle.hg38*-*Excludable.v2*. We also observed *hg38.Wold.hg38mitoExcludable* and *hg38.Lareau.hg38.full.Excludable* sets overlapping *hg38.Kundaje.GRCh38_unified_Excludable*, suggesting it contains NUMTs (Supplementary Figure S1D). Because of its agreement with other sets, we recommend *hg38.Kundaje.GRCh38_unified_Excludable* set and list other recommended sets Table 1.

## Discussion

Limited annotation remains the main problem when selecting exclusion sets as it remains unclear which method and/or data were used. Examples include Wold’s lab-generated “mitoblack” sets for mm9 and mm10 assemblies. Their curation method is unknown, and the exact number (123 regions), width distribution, and other characteristics suggest that one may be a liftOver version of the other. Similarly, it remains unknown why Bernstein’s lab-generated “Mint_Blacklist” hg19 and hg38 exclusion sets have a very large number of regions (9,035 and 12,052, respectively) as compared with under 1,000 regions for other exclusion sets. Additionally, hg19 and hg38 “full.blacklist” sets were generated by Caleb Lareau as a combination of NUMTs and unknown ENCODE exclusion sets, the source of which we were unable to infer. Given annotation shortcomings, we recommend using assembly-specific sets generated by a published method and, if relevant, combining them with other problematic region sets.

Most annotated exclusion sets were created via Blacklist, a tool for detecting regions with abnormally high signal and/or low mappability (*1*). These genomic properties are commonly accepted as problematic; however, they may not be exhaustive. The Peakpass algorithm was developed to learn genomic properties associated with problematic regions using a random forest model (*5*). It reported distance to nearest assembly gap or gene, and frequency of unique 4-mers or softmasked base pairs, as the most predictive of problematic regions. A limitation of Peakpass is that its extensive collection of Python, R, and bash scripts is poorly documented. A limitation of Blacklist, on the other hand, is computational resource requirements (64+ GB; CPU: 24+ cores, 3.4+ GHz/core) and disk storage (~ 1TB) due to a large number of required BAM files (hundreds). A recent preprint introduced the Greenscreen pipeline, a promising tool for identifying exclusion sets using as few as three ChIP-seq data. It reports a 99.9% overlap with a Blacklist-generated exclusion set, identical performance on ChIP-seq quality metrics but a smaller genome footprint (*13*). We utilized Blacklist as the most well-known tool to generate exclusion sets for the T2T-CHM13 and GRCm39 genome assemblies. The aforementioned tools detect problematic regions in ChIP-seq data; however, they may be different in data generated by other technologies due to different biochemical procedures (*14*). Additional collaborative efforts are needed to develop a consensus approach for defining well-documented exclusion sets.

## Acknowledgements

We thank Tim Triche and Stuart Lee for the helpful feedback and suggestions.

## Funding

This work was supported in part by the George and Lavinia Blick Research Scholarship to M.G.D., the Essential Open Source Software (EOSS) award from the Chan Zuckerberg Initiative (CZI) to M.I.L., the National Institutes of Health [R35-GM128645 to D.H.P.].

---

**Supplementary Figure S1.**
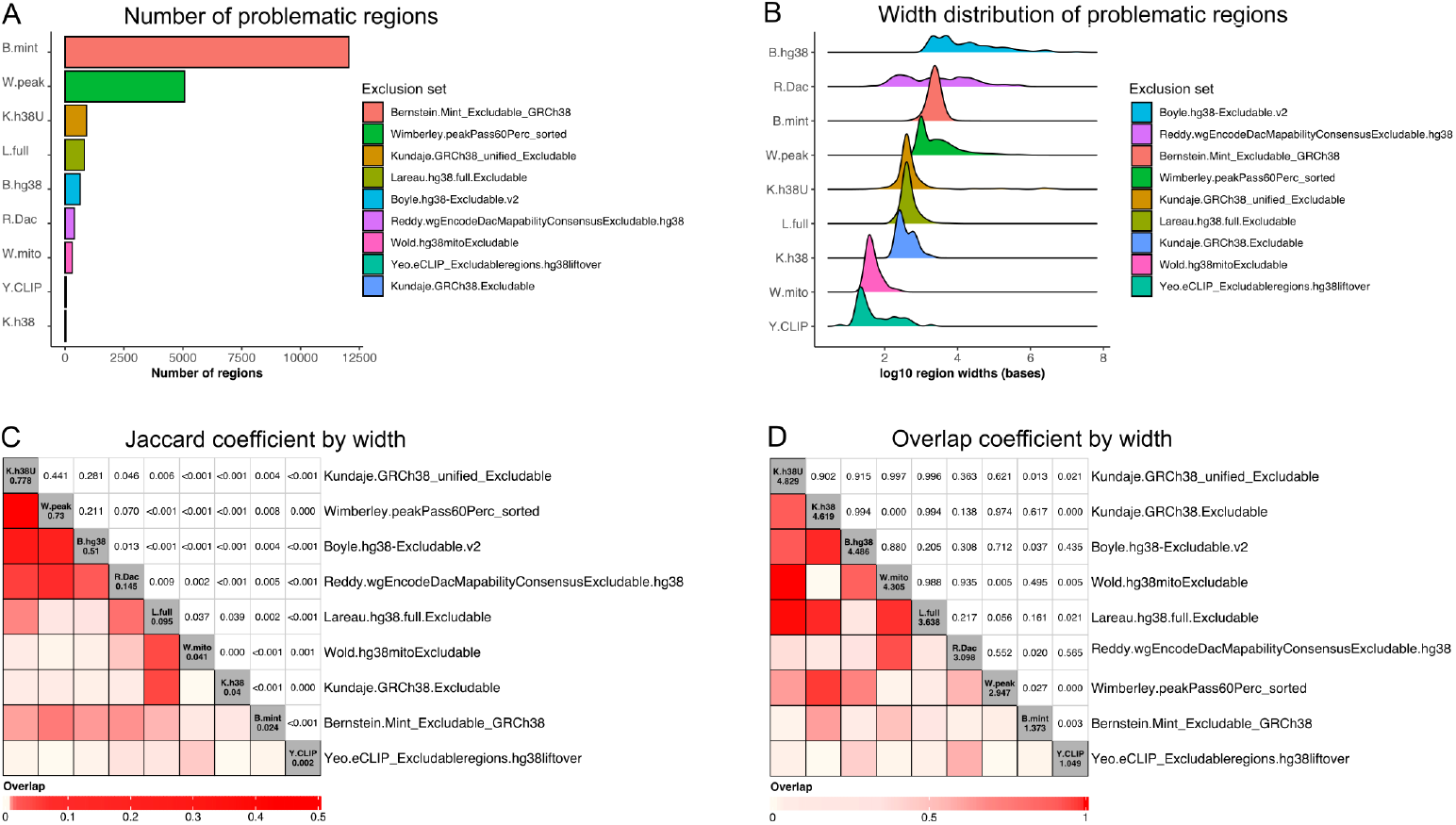
Characteristics of hg38 exclusion sets. (A) Number and (B) width distribution of problematic regions in hg38-specific exclusion sets. (C) Jaccard overlap 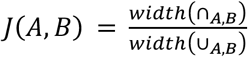 and (D) overlap coefficient 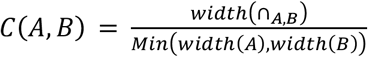 among hg38 exclusion sets by width. Diagonal counts represent sum of overlap coefficients of a list with all others.

**Supplementary Table S1.**
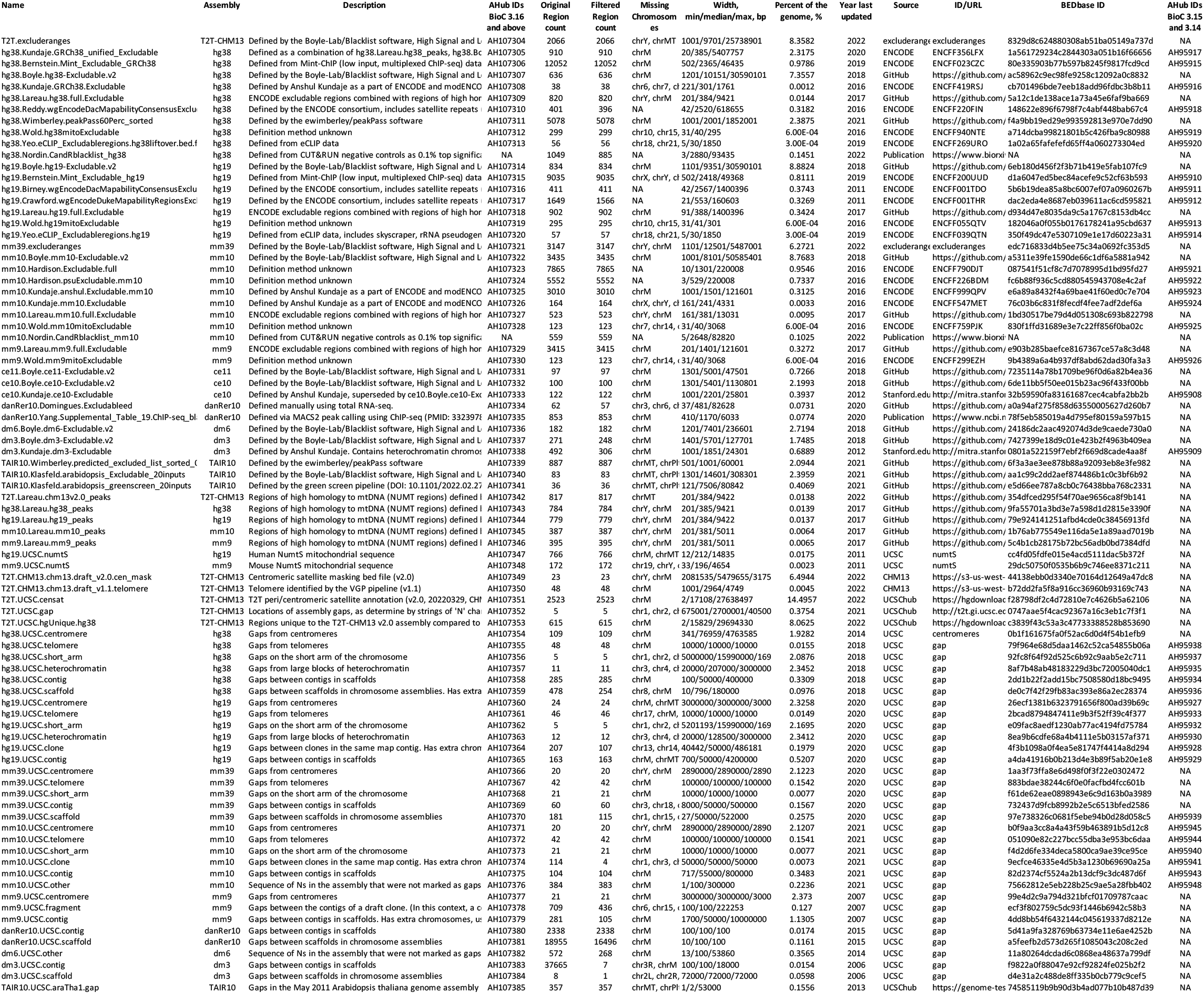
Characteristics of exclusion sets. “AHub IDs” - AnnotationHub IDs for objects in Bioconductor version 3.16 and above; “Original/Filtered regions” - the number of regions in the original set and in the subset to the assembled (autosomal) chromosomes; “ID/URL” - ENCODE ID or URL for data download. “BEDbase ID” - unique identifies for BEDbase.org API access, “Ahub IDs BioC 3.15 and 3.14” - AnnotationHub IDs for objects in Bioconductor version 3.14 and 3.15.

**Supplementary Table S2.** Gap overlap statistics for human and mouse exclusion sets. “% centromeres/short arms/telomeres covered” - proportion of gap regions covered by the corresponding exclusion set. “% regions intersecting centromeres/short arms/telomeres” - proportion of exclusion regions from a set covering gaps (number of overlapping regions over total).

| Name | %<br>centromeres<br>covered | % regions<br>intersecting<br>centromeres | % short<br>arms<br>covered | % regions<br>intersecting<br>short arms | %<br>telomeres<br>covered | % regions<br>intersecting<br>telomeres |
| --- | --- | --- | --- | --- | --- | --- |
| T2T.excluderanges | 93.58 | 2.27% (47/2066) | 74.13 | 1.79% (37/2066) | 94.59 | 2.18% (45/2066) |
| hg38.Bernstein.Mint_Excludable_GRCh38 | < 1.0 | 0.04% (5/12052) | 0 | 0% (0/12052) | 0 | 0% (0/12052) |
| hg38.Boyle.hg38-Excludable.v2 | 99.08 | 4.56% (29/636) | 58.91 | 0.47% (3/636) | 72.72 | 5.5% (35/636) |
| hg38.Kundaje.GRCh38_unified_Excludable | 99.4 | 2.97% (27/910) | 0 | 0% (0/910) | < 1.0 | 0.11% (1/910) |
| hg38.Kundaje.GRCh38.Excludable | < 1.0 | 7.89% (3/38) | 0 | 0% (0/38) | 0 | 0% (0/38) |
| hg38.Lareau.hg38.full.Excludable | < 1.0 | 0.37% (3/820) | 0 | 0% (0/820) | 0 | 0% (0/820) |
| hg38.Reddy.wgEncodeDacMapabilityConsensusExcludable.hg38 | 0 | 0% (0/396) | 0 | 0% (0/396) | < 1.0 | 0.25% (1/396) |
| hg38.Wimberley.peakPass60Perc_sorted | 59.6 | 26.98% (1370/5078) | < 1.0 | 0.08% (4/5078) | < 1.0 | 0.41% (21/5078) |
| hg38.Wold.hg38mitoExcludable | 0 | 0% (0/299) | 0 | 0% (0/299) | 0 | 0% (0/299) |
| hg38.Yeo.eCLIP_Excludableregions.hg38liftover | 0 | 0% (0/56) | 0 | 0% (0/56) | 0 | 0% (0/56) |
| hg38.Nordin.CandRblacklist_hg38 | 4.13 | 60.34% (534/885) | 0 | 0% (0/885) | 1.42 | 0.79% (7/885) |
| hg19.Bernstein.Mint_Excludable_hg19 | 0 | 0% (0/9035) | 0 | 0% (0/9035) | 0 | 0% (0/9035) |
| hg19.Birney.wgEncodeDacMapabilityConsensusExcludable | < 1.0 | 3.89% (16/411) | 0 | 0% (0/411) | < 1.0 | 0.24% (1/411) |
| hg19.Boyle.hg19-Excludable.v2 | 100 | 2.88% (24/834) | 92.26 | 0.48% (4/834) | 62.85 | 3.48% (29/834) |
| hg19.Crawford.wgEncodeDukeMapabilityRegionsExcludable | < 1.0 | 0.45% (7/1566) | 0 | 0% (0/1566) | 0 | 0% (0/1566) |
| hg19.Lareau.hg19.full.Excludable | < 1.0 | 1.77% (16/902) | 0 | 0% (0/902) | < 1.0 | 0.11% (1/902) |
| hg19.Wold.hg19mitoExcludable | 0 | 0% (0/295) | 0 | 0% (0/295) | 0 | 0% (0/295) |
| hg19.Yeo.eCLIP_Excludableregions.hg19 | 0 | 0% (0/57) | 0 | 0% (0/57) | 0 | 0% (0/57) |
| mm39.excluderanges | 83.3 | 1.11% (35/3147) | 43.66 | 0.89% (28/3147) | 76.19 | 0.51% (16/3147) |
| mm10.Boyle.mm10-Excludable.v2 | 88.07 | 1.08% (37/3435) | 100 | 0.12% (4/3435) | 76.19 | 0.47% (16/3435) |
| mm10.Hardison.Excludable.full | 0 | 0% (0/7865) | 0 | 0% (0/7865) | 0 | 0% (0/7865) |
| mm10.Hardison.psuExcludable.mm10 | 0 | 0% (0/5552) | 0 | 0% (0/5552) | 0 | 0% (0/5552) |
| mm10.Kundaje.anshul.Excludable.mm10 | 0 | 0% (0/3010) | 0 | 0% (0/3010) | 0 | 0% (0/3010) |
| mm10.Kundaje.mm10.Excludable | 0 | 0% (0/164) | 0 | 0% (0/164) | 0 | 0% (0/164) |
| mm10.Lareau.mm10.full.Excludable | 0 | 0% (0/523) | 0 | 0% (0/523) | 0 | 0% (0/523) |
| mm10.Wold.mm10mitoExcludable | 0 | 0% (0/123) | 0 | 0% (0/123) | 0 | 0% (0/123) |
| mm10.Nordin.CandRblacklist_mm10 | < 1.0 | 1.97% (11/559) | < 1.0 | 0.36% (2/559) | 0 | 0% (0/559) |

## Footnotes

1 https://docs.google.com/spreadsheets/d/1G4SkqUMiGcUlvR6homc7RW33nSOf4mS9QYJifsd4qo0

2 https://sites.google.com/site/anshulkundaje/projects/blacklists

3 https://www.encodeproject.org/search/?searchTerm=exclusion

4 https://github.com/caleblareau/mitoblacklist

5 https://github.com/marbl/CHM13

6 Example of BEDbase overview screen for hg38.Kundaje.GRCh38_unified_blacklist: http://bedbase.org/#/bedsplash/1a561729234c2844303a051b16f66656

7 Defined as a combination of *hg38.Lareau.hg38_peaks, hg38.Boyle.hg38-Excludable.v2*, and *hg38.Wimberley.peakPass60Perc_sorted*, followed by manual curation, https://www.encodeproject.org/files/ENCFF356LFX/

8 Defined by the PeakPass software, https://github.com/ewimberley/peakPass/raw/main/excludedlists/exclusion

## Notes

https://dozmorovlab.github.io/excluderanges/

### Competing Interest Statement

The authors have declared no competing interest.

